# The *Drosophila* microbiome has a limited influence on sleep, activity, and courtship behaviors

**DOI:** 10.1101/287045

**Authors:** Joel Selkrig, Farhan Mohammad, Soon Hwee Ng, Chua Jia Yi, Tayfun Tumkaya, Joses Ho, Yin Ning Chiang, Dirk Rieger, Sven Pettersson, Charlotte Helfrich-Förster, Joanne Y. Yew, Adam Claridge-Chang

## Abstract

In animals, commensal microbes modulate various physiological functions, including behavior. While microbiota exposure is required for normal behavior in mammals, it is not known how widely this dependency is present in other animal species. We proposed the hypothesis that the microbiome has a major influence on the behavior of the vinegar fly (*Drosophila melanogaster*), a major invertebrate model organism. Several assays were used to test the contribution of the microbiome on some well-characterized behaviors: defensive behavior, sleep, locomotion, and courtship in microbe-bearing, control flies and two generations of germ-free animals. None of the behaviors were largely influenced by the absence of a microbiome, and the small or moderate effects were not generalizable between replicates and/or generations. These results refute the hypothesis, indicating that the *Drosophila* microbiome does not have a major influence over several behaviors fundamental to the animal’s survival and reproduction. The impact of commensal microbes on animal behaviour may not be broadly conserved.

## Introduction

A human newborn is colonized during birth by diverse microbial species, initiating a complex and poorly understood molecular dialogue between the host and symbiotic microbes. Perturbation of this microbial community during early life is believed to disrupt a range of core physiological processes ^1–10^. A key physiological system found to be critically dependent on the early life microbiome is the brain; evidence shows that the microbiome affects brain function by modulating early brain development ^11–16^. For example, germ-free (GF) mice have abnormal function of their hypothalamic-pituitary-adrenal (HPA) axis, an important neurohormonal system; exposing GF mice to microbes during early life is sufficient to normalize HPA axis function ^15,17^. While microbial exposure in later life is insufficient to normalize the behavior of GF mice ^12,18^, normal function is restored by exposing either postnatal pups ^15,19^ or their mothers to microbial colonization ^16,20^. Thus, normal mammalian brain development and function rely on the microbiome. An intact mammalian gut microbiota is also required for normal anxiety-like behaviour and locomotion ^7,16,21^. However it is unclear whether this requirement is unique to mammals, or is a principle of brain development that applies broadly to other animal clades.

To investigate the microbe-brain relationship in other clades, we examined the role of the microbiome in the vinegar fly (*Drosophila melanogaster*), a major model organism for the neurobiology of behavior. *Drosophila* has several experimental advantages as a model for analyzing the microbiome-physiology interaction: it has a simple microbiome of typically only 5–30 taxa ^22,23^; is highly tractable for a range experimental manipulations; and offers large sample sizes that render high statistical precision. In addition, between mammals and flies, many of the fundamental mechanisms underlying host-microbial dynamics are conserved. Conserved microbe-interacting systems in the two clades include Toll-like receptor-based immune responses ^24^, metabolism ^25^, and insulin-like peptide signaling ^26,27^. Moreover, as with mammals, the *Drosophila* microbiome influences fly development across generations ^28,29^. While the microbiome clearly modulates *Drosophila* immunity and development ^27^, much less is known about commensal microbial effects on behavior.

We examined the role of the *Drosophila* microbiome in four well-characterized behaviors: anxiety-related wall following, locomotion, sleep, and courtship. In each of these behavioral paradigms, we found no substantial differences between GF and control flies. As the first analysis of cross-generational microbe-regulated *Drosophila* behaviour, this study indicates that the strong microbiome-brain interactions seen in mammals are not generalizable to all behaviors in all animal clades.

## Results

### Generation of germ-free adults

To test whether removal of the parental microbiome is capable of modulating behavioural outcomes in their offspring, we used *Wolbachia*-free Canton-S flies to generate first generation (F1) GF, second generation (F2) GF, and CV flies for behavioural profiling. We first homogenized and plated individual CV flies on MRS agar to characterize bacterial components in the wild-type *Drosophila* gut. PCR amplification and sequencing of 16S rRNA gene regions from individual colonies (*N* = 6) confirmed that previously identified bacterial genera *Lactobacillus* and *Acetobacter* were present in CV flies ^23,30^ (Table S1). To generate GF flies, fertilized Canton-S embryos were treated with bleach (Figure 1 A). F1 flies were deprived of microbes during embryogenesis whereas F2 flies were germ-free prior to fertilization. A nutrient-rich medium was chosen to reduce the magnitude of developmental delays as a potential confounding factor in observed phenotypes ^27^. The effectiveness of bacterial removal was confirmed by a negative PCR result for the bacterial 16S rRNA gene and the absence of colonies upon plating of fly homogenates onto agar permissive to the growth of gut bacteria (Figure 1 B). Consistent with previous reports in nutrient rich media, developmental delays (–1.33 days [95CI –1.47, –1.19]) were observed in GF flies, providing additional confirmation that the flies were indeed GF (Figure 1 C) ^27^.

**Figure 1.**
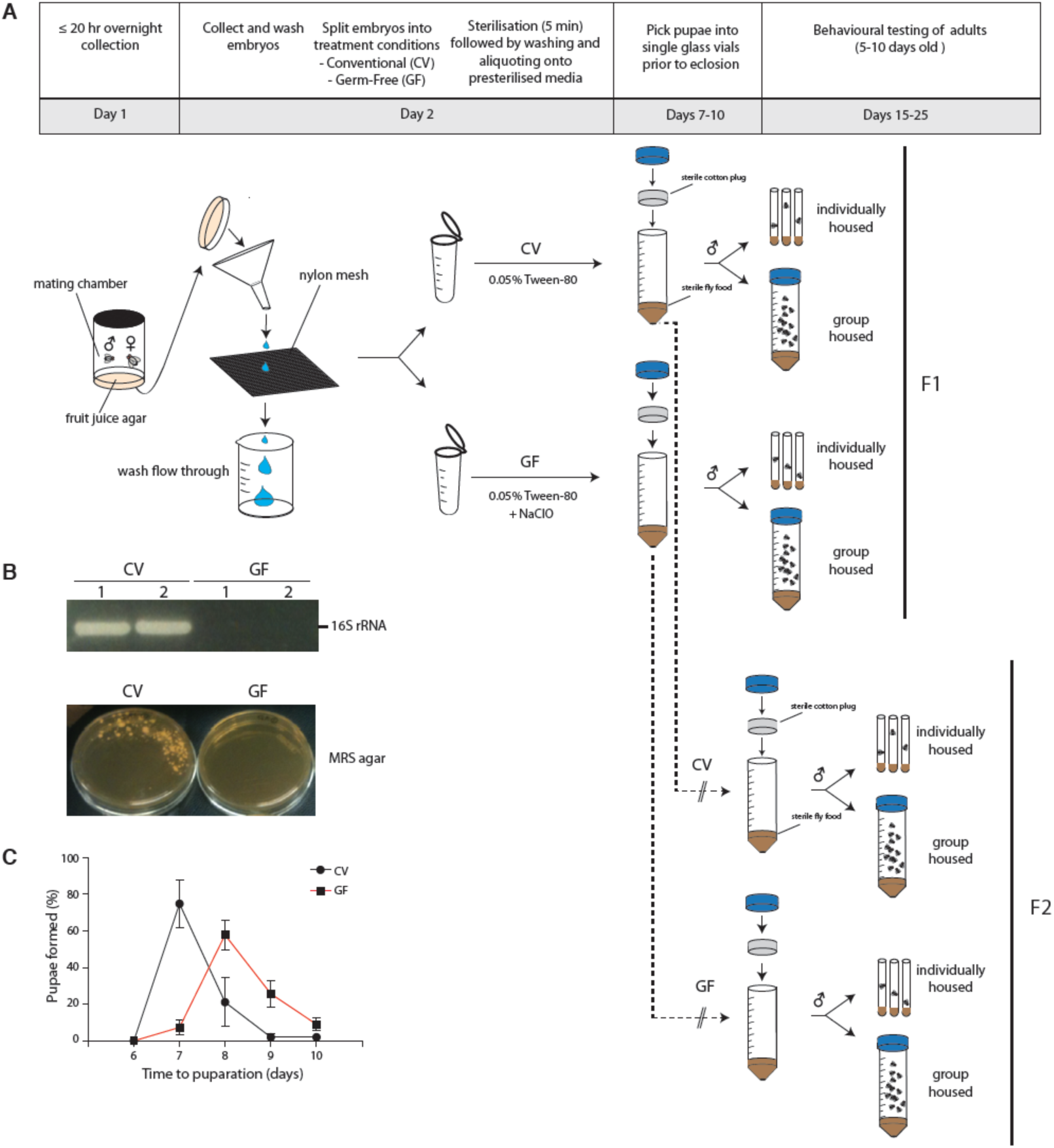
*Drosophila* embryo bleaching yields GF flies. **A.** Preparation of GF and CV flies prior to behavioural phenotyping. Prior to eclosion, pupae were picked into fresh vials and housed individually or in groups of 20. A full methodological description is provided in Methods. **B.** Validation of adult GF fly production by PCR of the 16s rRNA gene on fly homogenate (upper panel) in duplicate and culturing fly homogenate on MRS media (lower panel). A negative PCR result for the 16s rRNA gene and negative growth on MRS agar were used to confirm the absence of microbes. **C.** F1 GF flies were monitored for developmental delays by measuring the number of days it took for GF flies to form pupae. Error bars represent average % pupae formation +/– S.E.M. Bleaching of *Drosophila* embryos took 19% longer to form pupae post bleach treatment (−1.33 days [CI95 −1.47, −1.19], *P* = 1.0× 10^-4^, N = 112, 213].

### Individually housed GF and CV flies display similar wall-following behavior

The microbiome has been shown to influence rodent behaviour, including locomotion and anxiety ^7,12,14–16^. *Drosophila* wall following (WAFO) is governed by similar genetic mechanisms as rodent defense behaviours, and is an anxiety-related behaviour ^31^. We hypothesized that flies without a microbiota, like rodents, would display decreased defense behaviour; we also proposed that this low-anxiety phenotype would be counteracted by social isolation ^31^. We tested WAFO in two generations of GF and CV flies, raised in groups or isolation. Comparison of GF flies with control animals refuted this hypothesis: in three of four conditions, there were only trivial differences in WAFO between GF and CV flies (Figure 2 A-B, D). In the fourth condition, individually housed F2 GF flies exhibited a slight increase in WAFO relative to the CV controls (Figure 2 C), though the effect size was small (*g* = 0.495). Thus, the *Drosophila* microbiota does not have a major influence on anxiety-like behaviour.

**Figure 2.**
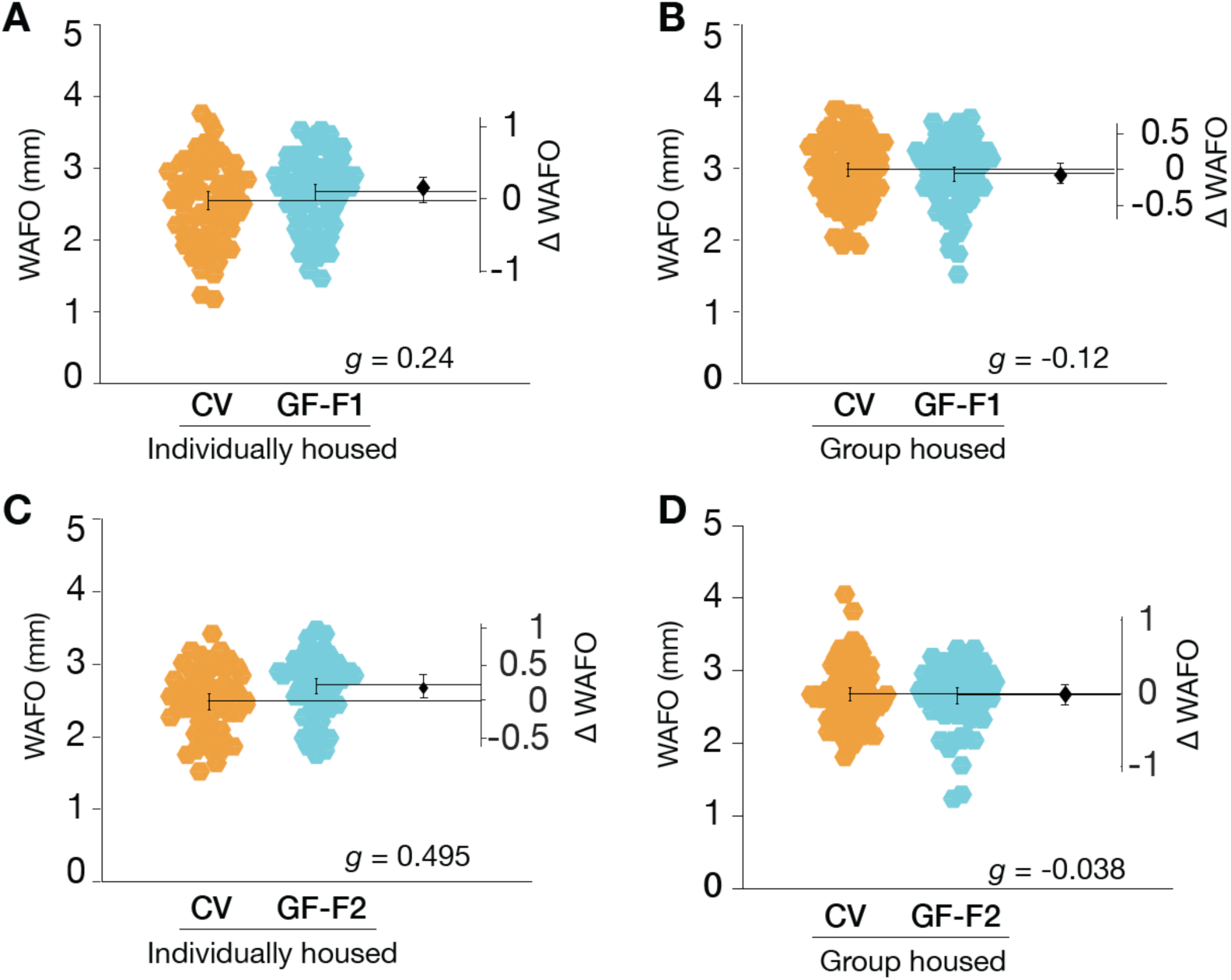
GF flies exhibit mostly trivial changes in wall following relative to CV controls. **A.** F1 individually housed GF flies did not exhibit substantially altered WAFO activity relative to CV counterparts (*g* = 0.236, *P* = 0.152, N = 75, 74). **B.** F1 group housed GF flies had largely unaltered WAFO activity (*g* = –0.118, *P* = 0.521, N = 96, 80). **C.** F2 individually housed exhibited a small elevation in WAFO activity (*g* = 0.495, *P* = 8.829e-03, N = 57, 56), whereas **D.** F2 group housed GF flies exhibited essentially no difference in WAFO activity from CV controls (*g* = –0.038, *P* = 5.125e-01, N = 89, 66).

### Locomotion is mildly elevated in second-generation GF flies

Tracking data from the WAFO assay was also analyzed to determine walking speed over a 10 min interval. In F1 adults, the removal of microbes had only trivial effects on locomotor activity; this was true for both socially naive and group housed GF flies (Figure 3 A, B). However, the F2 GF flies exhibited moderately elevated locomotor activity relative to CV controls (Figure 3 C, D); this was the case for both single- and grouped-housed animals. These data indicated that the microbiota plays no detectable role in modulating *Drosophila* brief-interval locomotor activity in the first generation, but suggested that the second generation were moderately hyperactive.

**Figure 3.**
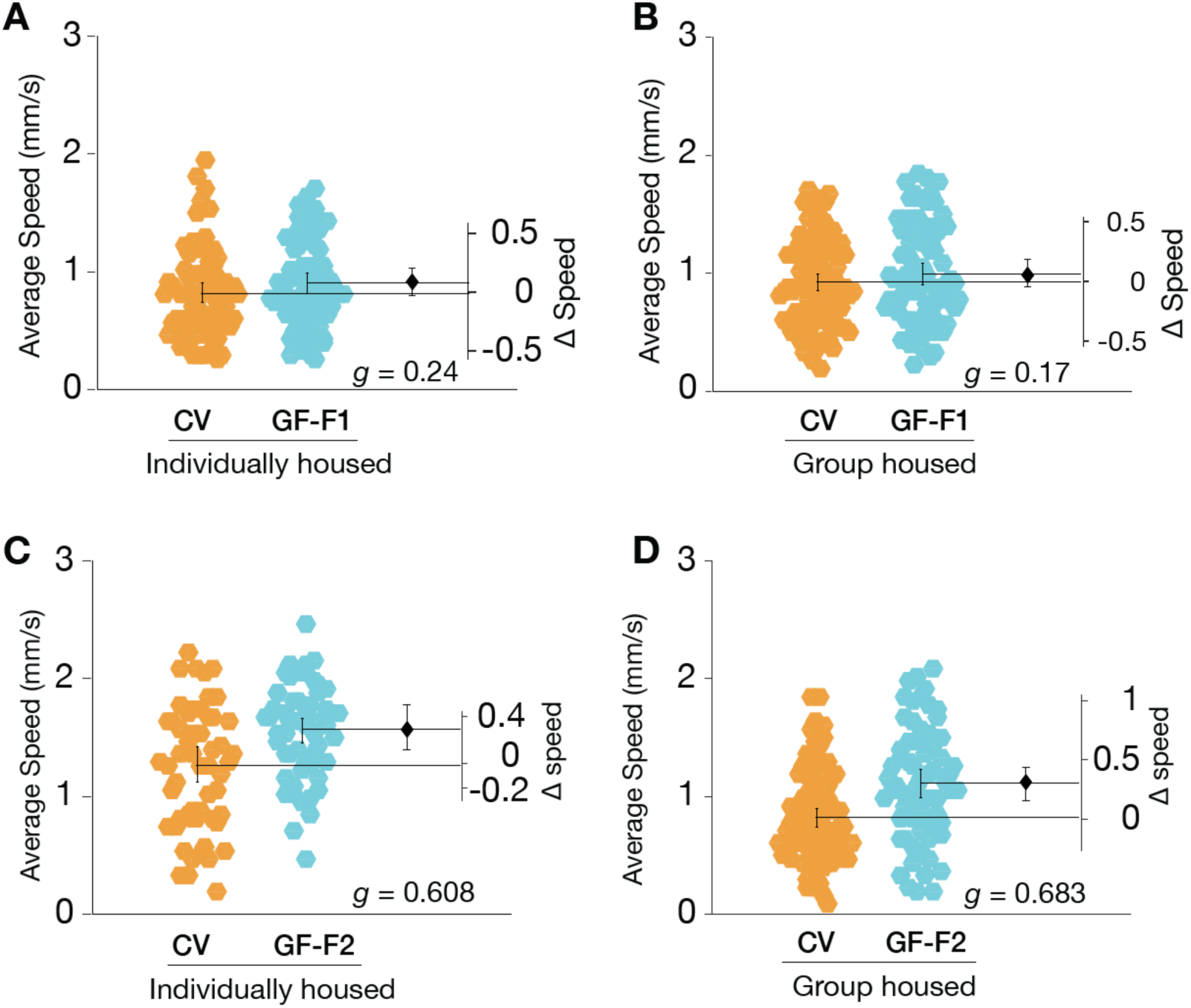
Removal of *Drosophila* microbiota mildly increases locomotion in the F2 generation. **A.** The average locomotion speed of F1 germ-free (GF) singly housed flies was elevated relative to conventional (CV) control animals (+0.09 mm/s [95CI −0.03, +0.21], *g* = 0.237, *P* = 1.002e-01, N = 75, 74). **B.** F1 group housed GF flies displayed an increase in locomotion relative to CV controls (+0.07 mm/s [95CI −0.05, +0.18], *g* = 0.17, *P* = 3.357e-01, N = 96, 80). **C.** Individually housed F2 GF flies walked 24.5 % faster than controls (+0.31 mm/s [95CI +0.15, +0.47], *g* = 0.608, *P* = 1.08 × 10^-3^, N = 57, 56). **D.** Group housed F2 GF flies walked 36.5 % faster than controls (+0.30 mm/s [95CI +0.18, +0.41], *g* = 0.683, *P* = 8.5 × 10^-5^, N = 89, 66).

### Microbe removal has little impact on activity or sleep

The increased activity observed in the 10 min WAFO assay led us to investigate hyperactivity in F2 GF flies. To examine this phenotype, we recorded activity over six days in both generations. Congruent with the WAFO speed data, the GF F1 flies had only a modest elevation in daytime activity (Figure 4 A–B); during the night, GF F1 flies were moderately less active than CV flies and slept slightly more (Figure 4 C–D). Concordant with the WAFO assay, F2 GF flies displayed moderately elevated daytime activity, along with a small increase in night time activity relative to controls (Figure 4 E–F); similarly, F2 GF flies slept less than controls (Figure 4 G–H).

**Figure 4.**
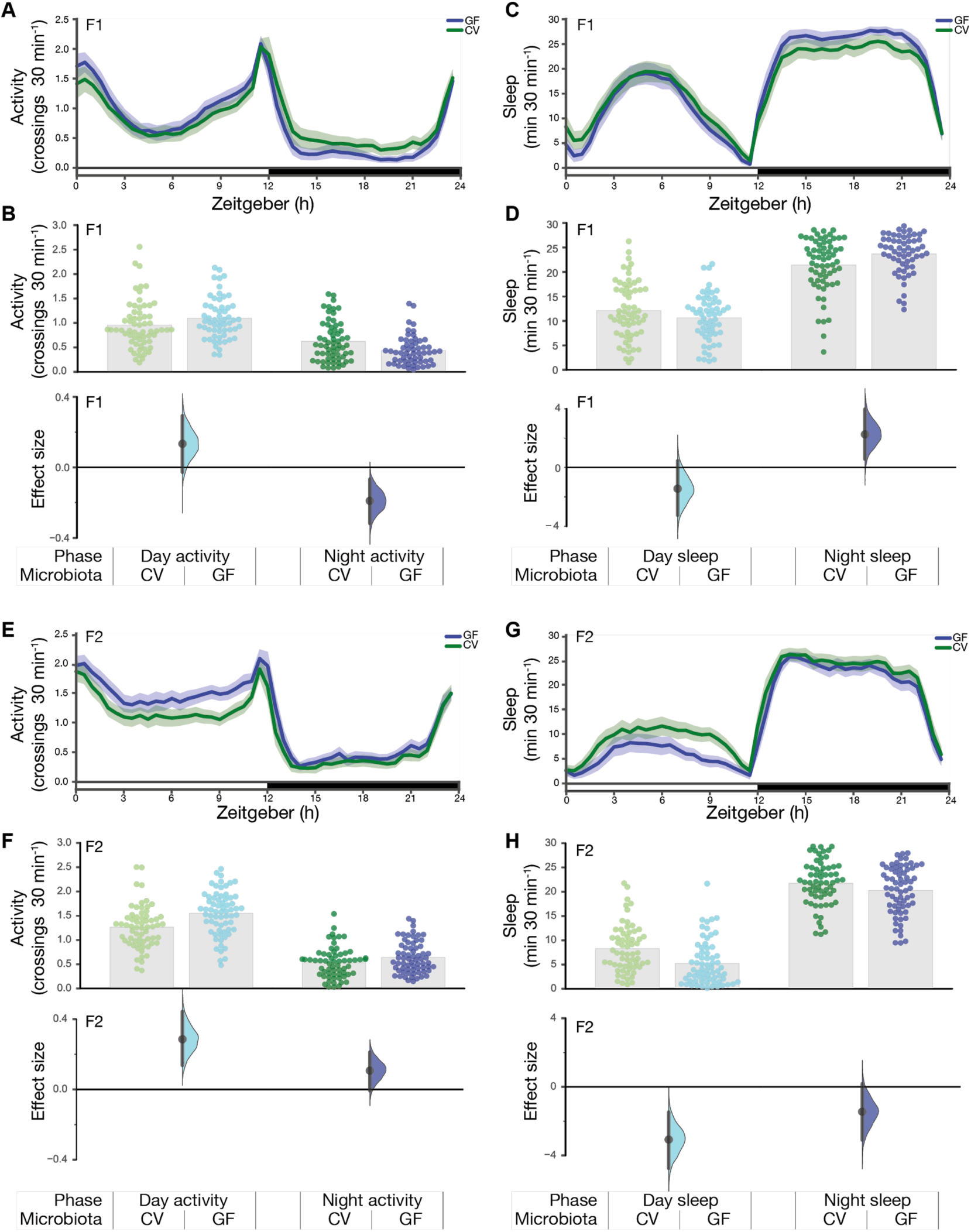
Germ-free flies do not have activity or sleep defects. **A.** Activity measured as photobeam crossings using the Drosophila Activity Monitoring System (DAMS) assay for group housed F1 CV and GF flies. **B.** In the day, GF flies made an average of 1.09 crossings/min (*N* = 59), while CV flies made an average of 0.96 crossings/min (*N* = 63), an increase of 0.13 [95CI −0.03, 0.29] (*P* = 0.11). At night, GF flies showed a slight decrease of –0.19 crossings/min [95CI −0.32, –0.07], *P* = 4.16 × 10^-3^ **C.** Sleep in group housed F1 CV and GF flies. **D.** In the day, GF flies slept for 10.61 minutes per 30 min (*N* = 59), while CV flies slept an average of 12.06/30 min (*N* = 63), a mean sleep decrease of −1.44 minutes [95CI −3.25, 0.46], *P* = 0.14. At night, GF flies showed a mild increase in sleep, with a mean difference of +2.26/30 min [95CI 0.56, 3.98], *P* = 0.01. **E.** Activity measured as beam crossings in the DAM assay for group housed F2 CV and GF flies. **F.** GF flies are slightly more active than CV flies. During the day, CV flies made an average of 1.26 crossings/min (*N* = 70), while GF flies made an average of 1.55 crossings/min (*N* = 67), an increase of +0.29 [95CI 0.14, 0.44] (*P* = 3.81 × 10^-4^). At night, GF flies showed an minor increase of +0.11 crossings/min [95CI –0.007, 0.21], *P* = 0.059. **G.** Sleep in group housed F2 CV and GF flies. **H.** During the day, GF flies slept for 5.23/30 min (*N* = 70), while CV flies slept an average of 8.30 min (*N* = 67), a mean sleep decrease of –3.07 minutes [95CI –4.62, –1.39], *P* = 3.08 × 10^-4^. At night, GF flies showed a mild decrease in sleep, with a mean difference of –1.45 min per 30 min [95CI – 3.09, 0.18], *P* = 0.089.

We aimed to generalize these findings to a second *Drosophila* stock, and to determine if the hyperactivity was reversible with microbiota reintroduction. However, in the replicate experiment, the elevated-activity phenotype was not reproduced. Indeed, relative to controls, the second batch of F2 GF flies were slightly less active (Figure 5 A–B). This difference between batches was also reflected in the sleep estimates (Figure 4 G–H, 5C–D). Additionally, microbiota reintroduction had almost no effect (Figure 5). To our knowledge, neither batch of experiments was flawed: sampling error and heterogeneity are typical causes of such differences between replicates ^32–34^. Thus, taken together, these results refute the hypothesis that the fly microbiota has a major influence on either *Drosophila* locomotor activity or sleep.

**Figure 5.**
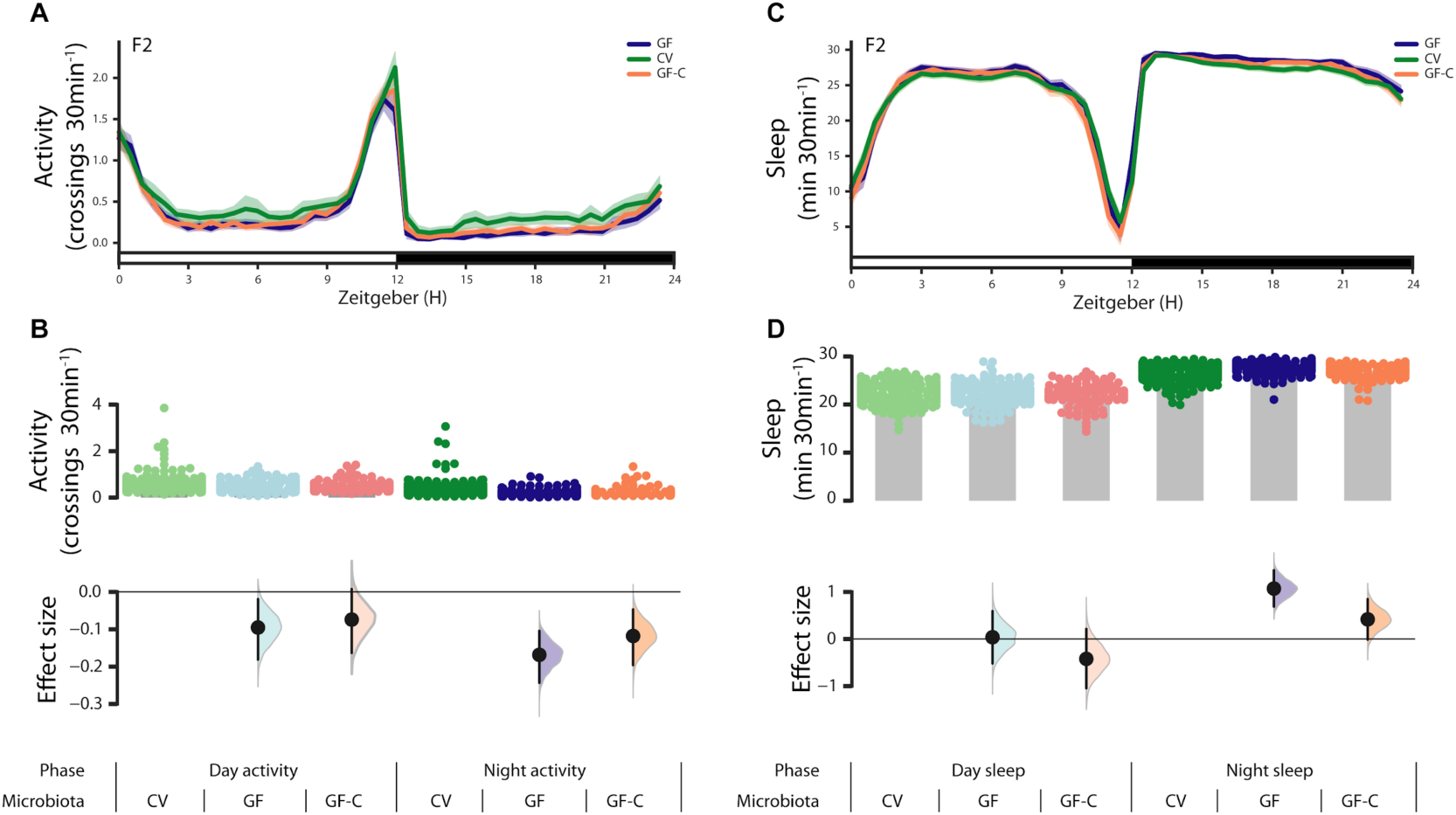
Conventional, germ-free, and colonized germ-free flies display similar sleep. **A.** Activity measured as photobeam crossings using the *Drosophila* Activity Monitoring System (DAMS) assay for group housed F2 CV, GF, and GF-C flies. **B.** In the day, CV, GF, and GF-C flies made an average of 0.60 crossings/min (*N* = 159), 0.51 crossings/min (*N* = 115), and 0.53 crossings/min (*N* = 90) respectively. Relative to CV, GF flies showed a decrease of −0.09 [95CI −0.18, −0.02] (*P* = 0.04). Relative to CV, GF-C flies showed a decrease of 0.07 [95CI −0.16, 0.01] (*P* = 0.15). At night, CV, GF, and GF-C flies made an average of 0.38 crossings/min (*N* = 159), 0.21 crossings/min (*N* = 115) and 0.27 crossings/min (*N* = 90) respectively. Relative to CV, GF flies showed a decrease of-0.17 [95CI −0.24, −0.10] (*P* = 0.00002). Relative to CV, GF-C flies showed a decrease of-0.12 [95CI −0.20, −0.05] (*P* = 0.009). **C.** Sleep estimates of group-housed F2 CV, GF, and GF-C flies. **D.** In the day, CV, GF, and GF-C flies slept for 22.30 minutes per 30 min (*N* = 159), 22.34 minutes per 30 min (*N* = 115), and 21.88 minutes per 30 min (*N* = 90) respectively. Relative to CV, GF flies showed a mean sleep increase of 0.04 minutes [95CI −0.52, 0.59], *P* = 0.9. Relative to CV, GF-C flies showed a mean sleep decrease of −0.42 minutes [95CI −1.05, 0.21], *P* = 0.17. At night, CV, GF, and GF-C flies slept for 26.54 minutes per 30 min (*N* = 159), 27.61 minutes per 30 min (*N* = 115) and 26.95 minutes per 30 min (*N* = 90) respectively. Relative to CV, GF flies showed a mean sleep increase of 1.07 minutes [95CI 0.69, 1.46], *P* = 0.0000004. Relative to CV, GF-C flies showed a mean sleep increase of 0.42 minutes [95CI −0.01, 0.85], *P* = 0.07.

### Conventional females are slightly more attractive than germ-free females

Previous reports have described microbiota-dependent mating preferences ^35,36^, though this is controversial ^37^. In these studies, dietary shifts were used to perturb microbiota compositions over multiple generations. In contrast, a recent report failed to detect a role for the *Drosophila* microbiota in mating preferences after dietary shifts and antibiotic exposure ^37^. To test whether direct microbe removal impacts microbiota-dependent attractiveness, conventional male flies were placed in a chamber together with decapitated GF and CV female bodies, and their courtship was monitored for 60 minutes (Figure 6 A). Courting wildtype males had a modest preference for CV females over GF females; this was true for both F1 (median proportion = 0.714 [95%CI 0.583; 0.855], *P* = 0.211) and F2 females (median proportion = 0.699 [95% CI 0.387; 0.945], *P* = 0.073). These results demonstrate that the *Drosophila* microbiota has just a minor effect on the attractiveness of either F1 or F2 females.

**Figure 6.**
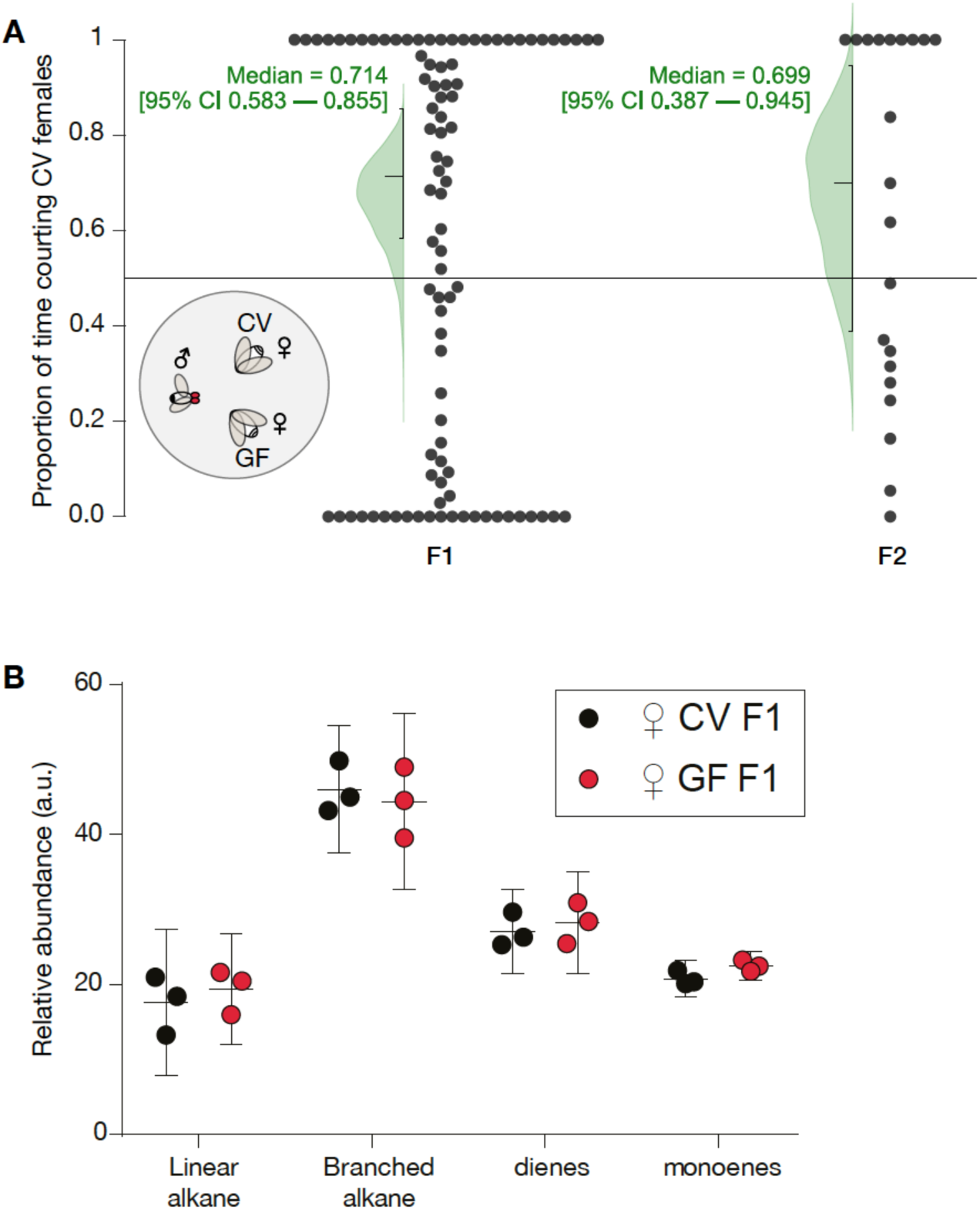
Germ-free female bodies are as attractive as conventional female bodies. **A. Inset:** A two-way choice courtship assay where conventionally raised (CR) males from fly stocks were aspirated into a chamber containing CV and GF female bodies. **Main axes:** 2-choice courtship assay of CV vs. GF female over a 60 min period. Only trials displaying courtship ≥ 1 min were analysed. Data was normalized by removing all non-courtship related behaviour throughout the 60 minute period. F1 (n = 92) and F2 (n = 21) courtships were analysed separately. The medians and their confidence intervals are given in green text; the green curves are bootstrap-estimated distributions of the medians. **B.** Relative abundance of CHC chemical classes determined by GC/MS. Data represent the average relative abundance (middle line) and confidence intervals (top and bottom lines) of 3 replicate sets per condition, with each set consisting of 8 females. Relative abundance (arbitrary units, a.u.) is calculated by dividing the area under each CHC peak byby the area under the hexacosane internal standard peak. Mean differences between CV and GF relative abundance: linear alkane −1.797 (95CI −8.681, +5.087, P = 0.9239), branched alkane +1.651 (95CI −5.232, +8.535, P = 0.9427), diene −1.174 (95CI −8.058, +5.71, P = 0.9830), monoene −1.71 (95CI −8.594, +5.173, P = 0.9355).

### GF females have a normal cuticular hydrocarbon profile

Female *Drosophila* attractiveness depends on the types of lipids on the cuticle ^38^. To see if there were differences that could explain the mild preference for CV females, we examined F1 female cuticular hydrocarbon (CHC) production with gas chromatography/ mass spectrometry (GC/MS). We detected only trivial differences in the overall amount of CHCs between CV and GF flies (Figure 6 B).

## Discussion

Here, we report the first examination of microbiota dependency behaviours in *Drosophila* across generations. We observed that microbiota removal had minor (and/or inconsistent) effects on anxiety-related behaviour ^7,16,20,39^, locomotion ^16,21^, sleep, and sexual attractiveness (Table 1). In summary, these findings suggest that the microbiota-gut-brain axis does not have a strong, consistent influence over several important *Drosophila* behaviours: defense, motor activity, and reproduction.

**Table 1.** Summary of effect sizes for all behavior assays. Summary of effect sizes for germ-free flies assayed for ^13^ behavioral metrics in ^26^ experiments. All effect sizes are Hedges’ *g*, except for courtship preference, which is a median proportion (md). Of the ^24^ Hedges’ *g* values, seven are moderate effect sizes; none are reproduced between replicate (Figure 5 vs. 4) or generation (F2 GF vs. F1 GF). IH = individually house; GH = group-housed; GF-C = colonized germ-free flies.

| Figure | Assay | F1 GF | F2 GF | F2 GF-C |
| --- | --- | --- | --- | --- |
| 2 | WAFO | 0.24 | 0.495 | ND |
| 2 | WAFO | -0.12 | -0.04 | ND |
| 3 | Locomotion | 0.24 | **0.61** | ND |
| 3 | Locomotion | 0.17 | **0.68** | ND |
| 4 | Activity - Day | 0.29 | **0.64** | ND |
| 4 | Activity - Night | **-0.53** | 0.33 | ND |
| 4 | Sleep - Day | -0.27 | **-0.65** | ND |
| 4 | Sleep -Night | 0.46 | -0.30 | ND |
| 5 | Activity - Day | ND | -0.26 | -0.19 |
| 5 | Activity - Night | ND | **-0.53** | -0.35 |
| 5 | Sleep - Day | ND | 0.02 | -0.18 |
| 5 | Sleep - Night | ND | **0.64** | 0.23 |
| 6 | Courtship | 0.71 md | 0.70 md | ND |

A range of studies have shown that germ-free rodents display abnormal defense behaviours, establishing that the indigenous mammalian microbiota is required for normal anxiety ^7,8,15,16,39,40^. Surprisingly, experiments in two generations of germ-free flies indicate that this interaction is either minimal or absent in lab-raised *Drosophila*. Similarly, while there is work showing that the microbiota affects rodent activity levels ^16^, the present study shows that this phenotype is not generalizable to flies. All the present experiments used sample sizes chosen to yield precise estimates of moderate effect sizes or larger (*N* = ≥56). If the microbiota influence on activity was moderate or large, it would have been measurable across generations and generalizable across conditions in a reproducible manner; this was not the case. Different stocks of the same laboratory strains can have differing microbiomes ^41,42^; while this could be used to explain the between-laboratories differences, it cannot explain the between-assays and between-generations discrepancies. Hence, we believe the parsimonious interpretation of the the varying, moderate-to-trivial effect sizes in the 10-min WAFO and 6-day DAMS locomotion assays is that microbiota role in fly behavioral activity is relatively minor.

Previous work relating the microbiota to *Drosophila* courtship used antibiotics or diet-induced shifts to assess its role in mating preferences ^35,36^. Our bleach-sterilization protocol removed the microbiota directly, allowing us to quantify the influence of the microbiota on female attractiveness. Our findings show that males court CV females with only a mild preference over GF females. This outcome contrasts with earlier reports demonstrating a major role for the microbiota in fly mate choice ^35,36^. The current observations are in line with recent work finding that diet-induced microbial changes did not alter mate choice ^37^.

Although many processes are conserved between mammals and flies, the present findings suggest the microbiota-gut-brain axis is not. There are at least three possible explanations for this difference. First, as rodent neuroscience uses small sample sizes and is affected by publication bias, these statistical factors may have amplified the rodent microbe–brain effect sizes that have been reported ^34,43,44^. Second, there are important differences between mammals and flies, for example, the brain’s relative size: it is plausible that the increased mammalian brains’ higher nutritional demand could mean that mammals rely more critically on microbial-derived nutrients ^45^. Third, it is worth considering that common lab-type fly strains have lost much of their original microbial diversity ^30^; it may be worthwhile examining how wild-type microbial strains influence (recently captive) wild-type fly behaviour ^42^.

In conclusion, although the microbiota impacts diverse aspects of *Drosophila* biology including immunity, metabolism, and development ^27,46–48^, the present evidence indicates that the microbiota influence on *Drosophila* behaviour is minor ^17,49^.

## Materials and Methods

### *Drosophila* husbandry

An isogenic Wolbachia-free *Drosophila melanogaster* Canton-S strain was used in all experiments. The absence of Wolbachia spp. was verified by PCR using wsp 81F (5’-TGGTCCAATAAGTGATGAAGAAAC-3’) and wsp 691R (5’-AAAAATTAAACGCTACTCCA-3’) ^50^ on crude DNA extract of pooled homogenate from five flies. Flies were raised on autoclaved cornmeal-dextrose-yeast agar food (<u>IMCB</u> and <u>EMBL</u> recipe available at Zenodo) at 25 °C under a 12:12 hr day/night cycle.

### Materials and media preparation

Bacteria isolated from *D. melanogaster* were cultivated on DeMan, Rogosa, and Sharpe (MRS) medium (Sigma cat. # 69966) containing 1.5% bacteriological agar. Embryos were sterilized using a 1:1 diluted 5% sodium hypochlorite (NaClO) bleach solution (FairPrice) and 0.05% Tween-80 (Sigma cat. # P8074). All solutions used were sterilized by autoclaving at 121 °C, 100 kPa for 15 minutes or passing through a 0.22 µm filter, where indicated, prior to use.

### Characterization of bacterial taxa in *Drosophila*

Bacterial colonies were isolated from the guts of *Drosophila* Canton-S on MRS agar and grown at 30 °C. Isolates were identified by sequencing the 16S rRNA PCR product generated using 8FE (5’-AGAGTTTGATCMTGGCTCAG-3’) and 1492R (5’-GGMTACCTTGTTACGACTT-3’) primers. PCR products were sequenced using the 8FE primer and were blasted against the National Center for Biotechnology Information database to assign bacterial identity, results are summarised in Table S1.

### Generation of germ-free and colonized *Drosophila*

F1 generation germ-free flies were prepared similarly to a previously described method ^51^. Briefly, eggs deposited overnight onto fruit juice agar were pooled then split into low protein binding 1.5 mL microfuge tubes with a ∼100 µL final egg bed volume. Subsequent steps were performed aseptically in a Class II biological safety cabinet. Eggs were washed twice with 1 mL of 0.05% Tween-80. Chorions were removed by exposing to 2.5% hypochlorite and 0.05% Tween-80 with gentle inverting of tubes for 5 min. Control conventional (CV) flies were prepared in parallel by exposure to 0.05% Tween-80 only. Eggs were washed twice with 1 mL of 0.05% Tween-80 solution followed by resuspension in 1 mL of 0.05% Tween-80 solution. 100 µL aliquots of treated eggs were then dispensed onto 10 mL of autoclaved fly food in 50 mL Falcon™ tubes. Pupae were isolated into autoclaved glass vials containing 2 mL fly food and capped with sterile cotton buds. Sexes were determined at eclosion and male experimental flies were either housed individually or in groups of 20. F2 generation GF and CV flies were prepared by transferring F1 adults to autoclaved food aseptically and F2 pupae were collected in the same way as F1. Quality control of GF fly preparation was performed at the end of each generation cycle with 5 adult flies. The absence of bacteria was verified by PCR for the 16S rRNA gene using 8FE (5’-AGAGTTTGATCMTGGCTCAG-3’) and 1492R (5’-GGMTACCTTGTTACGACTT-3’) and plating fly homogenates onto MRS agar ^47,51^.

For re-colonization of germ-free flies, monocultures of *Acetobacter pomorum* and *Lactobacillus* sp. were grown overnight at 30 °C with rotation in 2 mL of MRS broth. 50 µL aliquots of overnight culture from each strain was used to directly inoculate the surface of pre-autoclaved fly food inside 50 mL Falcon™ tubes. Dechorionated embryos were then deposited on top of the inoculated fly food and capped with sterilized cotton buds.

### Gas chromatography/mass spectrometry analysis

5 day old female *D. melanogaster* were anaesthetized on ice then collected in 1.8 mL glass vials (Wheaton, New Jersey, USA). 120 µL of hexane containing 10 µg/mL hexacosane (Sigma-Aldrich #241687) as an internal standard was added to triplicate vials (N_flies_ = 8 per vial) and incubated at room temperature for 20 minutes. 100 µL aliquots were transferred to fresh glass vials and were evaporated using a gentle stream of nitrogen. Samples were stored at –20 °C until analysis.

Analysis by gas chromatography/mass spectrometry (GC/MS) was performed with a QP2010 system (Shimadzu, Kyoto, Japan) equipped with a DB-5 column (5% Phenyl-methylpolysiloxane column; 30 m length, 0.25 mm ID, 0.25 µm film thickness; Agilent Technologies, CA, USA). Ionization was achieved by electron ionization (EI) at 70 eV. One microliter of the sample was injected using a splitless injector. The helium flow was set at 1.9 mL/min. The column temperature program began at 50 °C, increased to 210 °C at a rate of 35 °C /min, then increased to 280 °C at a rate of 3 °C/min. A mass spectrometer was set to unit mass resolution and 3 scans/ sec, from m/z 37 to 700. Chromatograms and mass spectra were analysed using GCMSsolution software (Shimadzu).

The relative abundance of each CHC species was calculated by dividing the area under the peak by the area of the hexacosane internal standard peak. The total area for each CHC chemical class (linear alkane, branched alkane, diene, or monoene) was calculated and the relative abundance from GF flies was then compared with that of the control CV flies using two-way ANOVA (GraphPad Prism 5, GraphPad Software Inc., CA, USA).

### Courtship choice

Courtship assays were performed using adult flies aged 7–10 days old, corresponding to sexual maturity. A socially naïve conventionally raised (CR) male was aspirated into a courtship chamber (35 × 10-mm) at 23.3 °C and 60% humidity containing one of each freshly decapitated GF female and a CV female 10-15 mm apart. Courtship behaviour (orienting, wing extension, wing vibration, or attempted copulation towards either female) was observed and recorded for 60 min. Courtship choice was calculated by taking a ratio of the amount of time spent by the male displaying courtship behaviours toward one female target to the total time the male spent courting either target. Trials with courtship activity lasting less than 1 min were omitted from analysis.

### Wall-following behaviour

The wall-following assay was performed as previously described ^31^. Briefly, for each behavioural assay, male flies were anesthetized in an ice-chilled vial for 30 seconds before being placed individually into an arena with forceps. The arena array was covered with an anti-reflection glass sheet (Edmund Optics, NJ, USA) and placed in an incubator. The array was lit from the sides by white LED microscope lamps (Falcon Illumination, Malaysia). To image the flies, an AVT Guppy F-046B CCD camera (Stemmer Imaging, Germany) equipped with a 12 mm CCTV-type lens was positioned above the behavioural arenas and connected to a computer via an IEEE 1394 cable. Flies were allowed to freely explore the arena during a 10-min test session. Flies’ positions were determined from a live video feed with CRITTA, custom software written in LabVIEW (National Instruments, U.S.A.), which extracts centroid x–y position data and records it to a binary file for offline analysis in MATLAB. Wall following (mean distance from the center of the chamber in mm) and mean speed (mm/sec) were calculated with custom scripts in MATLAB. The differences between control and experimental animal behaviors were reported as effect sizes (standardized mean differences, Hedges’ g). Hedges’ g and its bootstrapped 95% confidence interval were calculated with the Measurements of Effect Size toolbox (MES) in MATLAB ^52^. Hedges’ g is a variant of Cohen’s d standardized effect size that uses pooled SD and adjusts for bias ^53^.

### Activity and sleep assays

Group-housed male flies were placed in sterilized ^65^ mm glass tubes containing sterilized fly food on one end. The experiments were conducted at 12:12 hour light-dark cycle for a period of 6 days in an environment-controlled incubator (25 °C, 60% humidity). The first round of experiments were conducted at IMCB, the second at the Theodor-Boveri Institute. Fly activity was measured by recording the number of photobeam crossings in 1-minute bins using the *Drosophila* Activity Monitoring System (DAMS, Trikinetics, MA, USA). To generate the activity plots, six days of beam-crossing data were averaged into one 24-h interval and binned by 30 min. Then the binned activity data for the control and experimental flies were averaged separately, and represented as a line plot with the 95% confidence intervals. Sleep analysis was performed in Python using the pandas (McKinney 2011) and NumPy packages (Van Der Walt et al. 2011). Fly beam crossing data from the DAMS experiments were imported to the software and sleep events were identified as any period where the flies were inactive for at least 5 consecutive minutes (Shaw et al. 2000). Flies that did not move for 24 consecutive hours or more were considered dead and removed from the dataset. Then, the six-day sleep data was averaged into one day and binned by 30 minutes. The results were represented as a line plot along with the 95% confidence intervals. Mean sleep duration for the individual flies were represented with the bootstrapped 95% confidence intervals ^54^.

### Behavioral statistics and data analysis

For behavior data, estimation methods were used exclusively ^33^: no significance testing was conducted, and *P* values were reported *pro forma* only. Confidence intervals (95%) were calculated via bootstrap methods (10,000 samples taken) ^54^, and adjusted with bias correction and acceleration ^54^. A smoothed bootstrap was used for the median bootstrap calculations. The analysis was performed and visualized either in Python using the seaborn ^55,56^ and scikits-bootstrap packages ^57^. Several descriptors were used to describe effect size ranges: *trivial* (0 < *g* < 0.2); *small* (0.2 < *g* < 0.5); *moderate* (0.5 < *g* < 0.8); and *large* (*g* > 0.8) ^32^. Sample sizes are reported as the *N* of animals.

## Data availability

All data generated or analyzed during this study are included in this published article.

## Author Contributions

*Conceptualization*: JS and JYY; *Methodology*: JS (germ-removal methods); *Software*: JH and TT (Python) and FM (Matlab); *Investigation*: JS and SHN (germ removal, courtship), FM (locomotion, wall following), JYC (germ removal, sleep), and YNC (GC-MS analysis); *Data Analysis*: JH (courtship, sleep), TT (sleep), JYC (sleep) and FM (locomotion, WAFO); *Writing – Original Draft*: JS; *Writing – Revision*: JS, JYY, ACC, FM, SP; *Visualization*: JS, FM, JH, TT and JYY; *Supervision*: ACC and JYY; *Project Administration*: ACC and JYY; *Funding Acquisition*: ACC, SP and JYY.

## Acknowledgements

We thank Lisa Maier, Victoria Hewitt and Florian Szardenings for constructive criticism of this manuscript. We thank Shimadzu Asia Pacific for generously providing access to the GC-MS instrument.

