## Supplementary Materials for "The *Drosophila* microbiome has a limited influence on sleep, activity, and courtship behaviors"

**Table S1. Summary of bacterial isolates recovered from Canton S *Drosophila*.**
PCR amplification of the 16s rRNA genomic locus from individual bacterial colonies isolated from Canton S fly guts after growth on MRS agar. Sequences were generated using the 8FE-F primer.

| **Top BLASTn hit (Accession)** | **16s rRNA Sequence** |
|---|---|
| Uncultured *Lactobacillus* sp. gene for 16S ribosomal RNA, partial sequence, clone: wb-13 (LC036256.1) | AGTCGAACGAGCTGCGCCTAATGATAGTTGATGCTTGCATTAACTTGACTTAAGTTAGCA<br>GCGAGTGGCGAACTGGTGAGTAACACGTGGATAACCTGCCCAGAAGAAGGGGATAACACCTGGAAACAGATGCTAATACC<br>GTATAACAACGAAAACCACATGGTTTTCGTTTGAAAGATGGCCTTTGTGCTATCGCTTTTGGATGGATCCGCGGCGCATT<br>AGCTAGTTGGTGAGATAAAGGCTCACCAAGGCAATGATGCGTAGCCGACCTGAGAGGGTAATCGGCCACATTGGGACTGA<br>GACACGGCCCAGACTCCTACGGGAGGCAGCAGTAGGGAATCTTCCACAATGGACGAAAGTCTGATGGAGCAATGCCGCGT<br>GAGTGAAGAAGGGTTTCGGCTCGTAAAACTCTGTTGTTAGAGAAGAACGGGCGTGAGAGTAACTGCTCACGTCGCGACGG<br>TATCTAACCAGAAAGTCACGGCTAACTACGTGCCAGCAGCCGCGGTAATACGTAGGTGGCAAGCGTTGTCCGGATTTATT<br>GGGCGTAAAGCGAGCGCAGGCGGTCTTCTAAGTCTGATGTGAAAGCCTTCGGCTTAACCGGAGAAGTGCATCGGAAACTG<br>GGAAACTTGAGTGCAGAAGAGGACAGTGGAACTCCATGTGTAGCGGTGAAATGCGTAGATATATGGAGGAACACCAGTGG<br>CGAAGGCGGCTGTCTAGTCTGTAACTGACGCTGANGCTCGAAAGCATGGGTAGCAAACAGGATTAGATACCCTGGTAGTC<br>CATGCCGTAAACGATGAGTGCTAGGTGTTGGAGGGTTTCCGCCCTTCAGTGCCGCAGCTAACGCATTAAGCACTCCGCCT<br>GGGGAGTACNACCGCAAGGTTGAAACTCAAAGGAATTGACGGGGACCCGCACAAGCG |
| *Acetobacter pomorum* strain BDGP5 chromosome, complete genome (CP023657.1) | AGTCGCACGAAGGTTTCGGCCTTAGTGGCGGACGGGTGAGTAACGCGTAGGTATCT<br>ATCCATGGGTGGGGGATAACACTGGGAAACTGGTGCTAATACCGCATGACACCTGAGGGTCAAAGGCGTAAGTCGCCTGT<br>GGAGGAGCCTGCGTTTGATTAGCTAGTTGGTGGGGTAAAGGCCTACCAAGGCGATGATCAATAGCTGGTTTGAGAGGATG<br>ATCAGCCACACTGGGACTGAGACACGGCCCAGACTCCTACGGGAGGCAGCAGTGGGGAATATTGGACAATGGGGGCAACC<br>CTGATCCAGCAATGCCGCGTGTGTGAAGAAGGTCTTCGGATTGTAAAGCACTTTCGACGGGGACGATGATGACGGTACCC<br>GTAGAAGAAGCCCCGGCTAACTTCGTGCCAGCAGCCGCGGTAATACGAAGGGGGCTAGCGTTGCTCGGAATGACTGGGCG<br>TAAAGGGCGTGTAGGCGGTTTGTACAGTCAGATGTGAAATCCCCGGGCTTAACCTGGGAGCTGCATTTGATACGTGCAGA<br>CTAGAGTGTGAGAGAGGGTTGTGGAATTCCCAGTGTAGAGGTGAAATTCGTAGATATTGGGAAGAACACCGGTGGCGAAG<br>GCGGCAACCTGGCTCATTACTGACGCTGAGGCGCGAAAGCGTGGGGAGCAAACAGGATTAGATACCCTGGTAGTCCACGC<br>TGTAAACGATGTGTGCTAGATGTTGGGTGACTTAGTCATTCAGTGTCGCAGTTAACGCGTTAAGCACACCGCCTGGGGGA<br>GTACGGCCGCAAGGTTGAAACTCAAAGGAATTGACGGGGGCCCGCACAAGCG |
| *Acetobacter pomorum* strain BDGP5 chromosome, complete genome (CP023657.1) | AGTCGCACGAAGGTTTCGGCCTTAGTGGCGGACGGGTGAGTAACGCGTAGGTATCTATC<br>CATGGGTGGGGGATAACACTGGGAAACTGGTGCTAATACCGCATGACACCTGAGGGTCAAAGGCGTAAGTCGCCTGTGGA<br>GGAGCCTGCGTTTGATTAGCTAGTTGGTGGGGTAAAGGCCTACCAAGGCGATGATCAATAGCTGGTTTGAGAGGATGATC<br>AGCCACACTGGGACTGAGACACGGCCCAGACTCCTACGGGAGGCAGCAGTGGGGAATATTGGACAATGGGGGCAACCCTG<br>ATCCAGCAATGCCGCGTGTGTGAAGAAGGTCTTCGGATTGTAAAGCACTTTCGACGGGGACGATGATGACGGTACCCGTA<br>GAAGAAGCCCCGGCTAACTTCGTGCCAGCAGCCGCGGTAATACGAAGGGGGCTAGCGTTGCTCGGAATGACTGGGCGTAA<br>AGGGCGTGTAGGCGGTTTGTACAGTCAGATGTGAAATCCCCGGGCTTAACCTGGGAGCTGCATTTGATACGTGCAGACTA<br>GAGTGTGAGAGAGGGTTGTGGAATTCCCAGTGTAGAGGTGAAATTCGTAGATATTGGGAAGAACACCGGTGGCGAAGGCG<br>GCAACCTGGCTCATTACTGACGCTGAGGCGCGAAAGCGTGGGGAGCAAACAGGATTAGATACCCTGGTAGTCCACGCTGT<br>AAACGATGTGTGCTAGATGTTGGGTGACTTAGTCATTCAGTGTCGCAGTTAACGCGTTAAGCACACCGCCTGGGGAGTAC<br>GGCCGCAAGGTTGAAACTCAAAGGAATTGACGGGGGCCCGCACAAGCG |
| *Acetobacter pomorum* strain BDGP5 chromosome, complete genome (CP023657.1) | AGTCGCACGAAGGTTTCGGCCTTAGTGGCGGACGGGTGAGTAACGCGTANNNNNNNNN<br>CCNTGGGTGGGGGATAACACTGGGAAACTGGTGCTAATACCGCATGACACCTGAGGGTCAAAGGCGTAAGTCGCCTGTGG<br>AGGAGCCTGCGTTTGATTAGCTAGTTGGTGGGGTAAAGGCCTACCAAGGCGATGATCAATAGCTGGTTTGAGAGGATGAT<br>CAGCCACACTGGGACTGAGACACGGCCCAGACTCCTACGGGAGGCAGCAGTGGGGAATATTGGACAATGGGGGCAACCCT<br>GATCCAGCAATGCCGCGTGTGTGAAGAAGGTCTTCGGATTGTAAAGCACTTTCGACGGGGACGATGATGACGGTACCCGT<br>AGAAGAAGCCCCGGCTAACTTCGTGCCAGCAGCCGCGGTAATACGAAGGGGGCTAGCGTTGCTCGGAATGACTGGGCGTA<br>AAGGGCGTGTAGGCGGTTTGTACAGTCAGATGTGAAATCCCCGGGCTTAACCTGGGAGCTGCATTTGATACGTGCAGACT<br>AGAGTGTGAGAGAGGGTTGTGGAATTCCCAGTGTAGAGGTGAAATTCGTAGATATTGGGAAGAACACCGGTGGCGAAGGC<br>GGCAACCTGGCTCATTACTGACGCTGAGGCGCGAAAGCGTGGGGAGCAAACAGGATTAGATACCCTGGTAGTCCACGCTG<br>TAAACGATGTGTGCTAGATGTTGGGTGACTTAGTCATTCAGTGTCGCAGTTAACGCGTTAAGCACACCGCCTGGGGAGTA<br>CGGCCGCAAGGTTGAAACTCAAAGGAATTGACGGGGGCCCGCACAAGCG |
| Uncultured *Lactobacillus* sp. gene for 16S ribosomal RNA, partial sequence, clone: wb-13 (LC036256.1) | AGTCGAACGAGCTGCGCCTAATGATAGTTGATGCTTGCATTAACTTGACTTAAGTTAGCA<br>GCGAGTGGCGAACTGGTGAGTAACACGTGGATAACCTGCCCAGAAGAAGGGGATAACACCTGGAAACAGATGCTAATACC<br>GTATAACAACGAAAACCACATGGTTTTCGTTTGAAAGATGGCCTTTGTGCTATCGCTTTTGGATGGATCCGCGGCGCATT<br>AGCTAGTTGGTGAGATAAAGGCTCACCAAGGCAATGATGCGTAGCCGACCTGAGAGGGTAATCGGCCACATTGGGACTGA<br>GACACGGCCCAGACTCCTACGGGAGGCAGCAGTAGGGAATCTTCCACAATGGACGAAAGTCTGATGGAGCAATGCCGCGT<br>GAGTGAAGAAGGGTTTCGGCTCGTAAAACTCTGTTGTTAGAGAAGAACGGGCGTGAGAGTAACTGCTCACGTCGCGACGG<br>TATCTAACCAGAAAGTCACGGCTAACTACGTGCCAGCAGCCGCGGTAATACGTAGGTGGCAAGCGTTGTCCGGATTTATT<br>GGGCGTAAAGCGAGCGCAGGCGGTCTTCTAAGTCTGATGTGAAAGCCTTCGGCTTAACCGGAGAAGTGCATCGGAAACTG<br>GGAAACTTGAGTGCAGAAGAGGACAGTGGAACTCCATGTGTAGCGGTGAAATGCGTAGATATATGGAGGAACACCAGTGG<br>CGAAGGCGGCTGTCTAGTCTGTAACTGACGCTGANGCTCGAAAGCATGGGTAGCAAACAGGANTAGATACCCTGGTAGTC<br>CATGCCGTAAACGATGAGTGCTAGGTGTTGGAGGGTTTCCGCCCTTCAGTGCCGCAGCTAACGCATTAAGCACTCCGCCT<br>GGGGAGTACGACCGCAAGGTTGAAACTCAAAGGAATTGACGGGGACCCGCACAAGCG |
| Uncultured *Lactobacillus* sp. gene for 16S ribosomal RNA, partial sequence, clone: wb-13 (LC036256.1) | AGTCGAACGAGCTGCGCCTAATGATAGTTGATGCTTGCATTAACTTGACTTAAGTTAGC<br>AGCGAGTGGCGAACTGGTGAGTAACACGTGGATAACCTGCCCAGAAGAAGGGGATAACACCTGGAAACAGATGCTAATAC<br>CGTATAACAACGAAAACCACATGGTTTTCGTTTGAAAGATGGCCTTTGTGCTATCGCTTTTGGATGGATCCGCGGCGCAT<br>TAGCTAGTTGGTGAGATAAAGGCTCACCAAGGCAATGATGCGTAGCCGACCTGAGAGGGTAATCGGCCACATTGGGACTG<br>AGACACGGCCCAGACTCCTACGGGAGGCAGCAGTAGGGAATCTTCCACAATGGACGAAAGTCTGATGGAGCAATGCCGCG<br>TGAGTGAAGAAGGGTTTCGGCTCGTAAAACTCTGTTGTTAGAGAAGAACGGGCGTGAGAGTAACTGCTCACGTCGCGACG<br>GTATCTAACCAGAAAGTCACGGCTAACTACGTGCCAGCAGCCGCGGTAATACGTAGGTGGCAAGCGTTGTCCGGATTTAT<br>TGGGCGTAAAGCGAGCGCAGGCGGTCTTCTAAGTCTGATGTGAAAGCCTTCGGCTTAACCGGAGAAGTGCATCGGAAACT<br>GGGAAACTTGAGTGCAGAAGAGGACAGTGGAACTCCATGTGTAGCGGTGAAATGCGTAGATATATGGAGGAACACCAGTG<br>GCGAAGGCGGCTGTCTAGTCTGTAACTGACGCTGANGCTCGAAAGCATGGGTAGCAAACAGGATTAGATACCCTGGTAGT<br>CCATGCCGTAAACGATGAGTGCTAGGTGTTGGAGGGTTTCCGCCCTTCAGTGCCGCAGCTAACGCATTAAGCACTCCGCC<br>TGGGGAGTACGACCGCAAGGTTGAAACTCAAAGGAATTGACGGGGACCCGCACAAGCG |
